# Global brain maintenance predicts well-preserved cognitive function: A pooled analysis of three longitudinal population-based Swedish cohorts

**DOI:** 10.1101/2025.03.04.641447

**Authors:** Saana M. Korkki, Justinas Narbutas, Alireza Salami, Lina Rydén, Grégoria Kalpouzos, Micael Andersson, Silke Kern, Johan Skoog, Eric Westman, Ingmar Skoog, Debora Rizzuto, Lars Nyberg

## Abstract

Substantial heterogeneity in cognitive ageing is well documented. Such heterogeneity has been attributed to individual differences in *brain maintenance –* i.e., the relative preservation of neural resources in ageing. However, large-scale longitudinal evidence is lacking. We pooled data from three population-based Swedish cohorts (Betula, *N* = 196; SNAC-K, *N* = 472; H70, *N* = 688; aged 60-93 years at baseline, follow-up duration up to 7 years) to assess whether global brain maintenance is associated with better preserved cognition in ageing, and to identify lifestyle predictors of brain maintenance. In each cohort, global brain integrity was indexed by the volume of the lateral ventricles (adjusted for total intracranial volume), and general cognitive function based on a principal component analysis of four age-sensitive cognitive domains. Participants were classified into subgroups of low (i.e., ‘aged’) versus high (i.e., ‘youth-like’) brain integrity based on ventricular volume estimates available for a younger reference sample in one of the cohorts (Betula, 25-55 years, *N* = 60). Subgroup differences in cognition at baseline and over the follow-up were assessed with ANCOVAs and linear mixed effects models. Logistic regressions were used to examine lifestyle predictors of brain maintenance. Across cohorts, 881 individuals (64.97%) were classified into the high brain integrity subgroup at baseline and 409 individuals (49.82%) over the follow-up. Maintenance of more youth-like brain integrity was associated with better baseline cognition (*p* < .001) and less cognitive decline longitudinally (*p* < .001). Moreover, lower cardiovascular disease (CVD) risk and the absence of diabetes predicted brain maintenance at baseline (CVD risk, OR = 0.80, 95% CI [0.68, 0.93]; diabetes, OR = 0.39, 95% CI [0.26, 0.59]) and over the follow-up (CVD risk, OR = 0.79, 95% CI [0.64, 0.96]; diabetes, OR = 0.53, 95% CI [0.29, 0.94]). These findings underscore brain maintenance as a key determinant of cognitive ageing and highlight the importance of managing cardiovascular and metabolic disease risk factors for promotion of brain and cognitive health in later life.

## Introduction

Despite mean-level cognitive decline, there is substantial heterogeneity in cognitive ageing trajectories, with some older individuals exhibiting levels of function similar to younger counterparts^1,2^. In terms of the neural substrates, this heterogeneity has been attributed to individual differences in *reserve* (i.e., greater resilience to deterioration due to surplus neural resources), *maintenance* (i.e., less deterioration of neural resources), and *compensation* (i.e., recruitment of additional neural resources)^3^. Notably, the account of *brain maintenance* suggests that individuals who display less age-related neural decline also exhibit better preserved cognition in later life^1^. Previous neuroimaging studies have provided evidence for both general and specific forms of brain maintenance^4,5^. For instance, Cox and colleagues^4^ observed overall cortical atrophy to account for longitudinal declines in both general- and domain-specific cognitive function, whereas others have indicated additional contributions of, for example, hippocampal maintenance to episodic memory^5^. However, much of the existing evidence stems from single cohort studies, and large-scale longitudinal investigations of the role of brain maintenance in cognitive ageing are currently lacking.

In terms of structural brain changes, advancing adult age is associated with decreases in cortical and subcortical grey matter volume^6–8^, reduced integrity of white matter tracts connecting different brain regions^9^, and increases in ventricular volume^6–8^. Ventricular volume remains relatively stable until mid-life and displays accelerated increases thereafter^6,10^, predicting cognitive decline in healthy older adults^11^, and serving as a marker for cognitive impairment and dementia^12^. Previous studies have demonstrated higher reliability of ventricular volume estimates in comparison to some other subcortical and cortical structures^13,14^, and good reproducibility across varying scanners and acquisition sequences^15–17^, making it a suitable metric of global brain integrity for multi-cohort investigations of cognitive ageing.

Moreover, the determinants of individual differences in brain maintenance remain poorly understood, likely encompassing both genetic and environmental factors^18^. Evidence from population-based cohorts and intervention studies suggests an important role for midlife and old age lifestyle factors in the maintenance of brain health in ageing^19–22^. In particular, higher burden of cardiovascular disease risk factors (e.g., hypertension, physical inactivity, diabetes) has previously been linked to cognitive decline^23^ and dementia^24^, as well as poorer outcomes on various markers of brain structural integrity^25–27^. Additionally, later life differences in brain and cognition may be affected by environmental influences during early life periods^28^. Recently, secular trends for brain structural measures have been reported, with later-born cohorts displaying higher brain integrity^29^, aligning with previous observations of birth cohort effects on cognitive performance^30^.

In this study, we pooled structural magnetic resonance imaging (MRI) data from three population-based Swedish cohorts (Betula, SNAC-K, H70) from the National E-Infrastructure for Aging Research (NEAR) database to investigate the role of global brain maintenance in cognitive ageing and to identify potential lifestyle predictors of brain maintenance. Leveraging estimates of ventricular volume available for a younger reference sample in one of the cohorts (Betula), we classified each older adult into high (i.e., ‘youth-like’) or low (i.e., ‘aged’) brain integrity subgroup and assessed their differences in general cognitive function at baseline and over a follow-up period of up to 7 years. We expected maintenance of global brain integrity to be associated with better preserved cognitive function, and to be linked to a lower burden of cardiovascular disease risk factors previously implicated in brain health^22^. Lastly, we examined potential birth cohort effects on global brain integrity in 70-72-year-old individuals born up to 15 years apart across the three cohorts, with the prediction that later born cohorts would display better brain integrity^29^.

## Methods

### Participants

Data were drawn from three population-based longitudinal Swedish cohorts (Betula, SNAC-K, and H70) from the NEAR database (https://www.near-aging.se/). The study protocols and recruitment procedures for each cohort have been described in detail previously^31–34^. The cohorts located in three different Swedish cities; Betula in Umeå, SNAC-K in Stockholm, and H70 in Gothenburg. Baseline age range was 60-93 years for SNAC-K, 70-72 years for H70, and 25-81 years for Betula. Data from young to middle-aged adults (25-55 years, *N* = 60) in Betula was used for estimation of a ventricular volume threshold based on which older individuals in each cohort were classified into subgroups of high (i.e., ‘youth-like’) versus low (i.e., ‘aged’) brain integrity (see Statistical analyses). All other analyses were restricted to the older adults in Betula (≥ 60 years).

Individuals with a diagnosis of dementia, indication of other neurological disease or condition (e.g., stroke, Parkinson’s disease), or a low baseline score (< 24) on the Mini-Mental State Examination (MMSE^35^) were excluded from the analyses. The baseline magnetic resonance imaging (MRI) sample consisted of 196 individuals for Betula, 472 individuals for SNAC-K, and 688 individuals for H70. For longitudinal analyses, participants with at least one follow-up datapoint were included, resulting in a longitudinal MRI sample of 120 individuals for Betula, 294 individuals for SNAC-K, and 407 individuals for H70. For Betula, data was available for a maximum of two follow-ups occurring 4 and 7 years after the baseline. For SNAC-K, data was available for a maximum of two follow-ups that occurred 3 and 6 years after the baseline, with all participants invited for the 6-year follow-up, and only individuals ≥ 78 years old at baseline for the 3-year follow-up. For H70, one follow-up that occurred 5 years after the baseline was available. Baseline characteristics of older participants in each cohort are reported in Table 1 (see Supplementary Table S1 for characteristics of Betula younger adults).

**Table 1.** Baseline characteristics of older participants (≥ 60 years) in each cohort.

|  | BETULA |  | SNAC-K |  | H70 |  |
| --- | --- | --- | --- | --- | --- | --- |
|  | N | Mean (SD) /<br>Percentage | N | Mean (SD) /<br>Percentage | N | Mean (SD) /<br>Percentage |
| Age<br>(years) | 196 | 69.04<br>(6.37) | 472 | 70.26<br>(8.99) | 688 | 70.23<br>(0.44) |
| Sex | 196 | 48.98% F | 472 | 58.69% F | 688 | 53.34% F |
| Education<br>(years) | 194 | 12.52<br>(4.25) | 470 | 12.69<br>(4.06) | 687 | 13.40<br>(4.14) |
| MMSE | 196 | 28.04<br>(1.48) | 472 | 29.14<br>(1.04) | 687 | 29.09<br>(1.17) |
| Ventricular<br>volume | 196 | 34758.13<br>(17911.12) | 472 | 33066.92<br>(18246.29) | 688 | 33312.33<br>(15824.02) |
| ICV | 196 | 1479421.19<br>(172912.11) | 472 | 1515201.88<br>(246437.03) | 688 | 1502199.68<br>(159111.56) |
| BMI | 196 | 26.45<br>(3.57) | 459 | 26.10<br>(3.75) | 687 | 25.90<br>(4.29) |
| BP systolic | 196 | 142.30<br>(17.61) | 455 | 142.60<br>(19.21) | 686 | 139.84<br>(18.70) |
| Smoker | 196 | 8.16% | 472 | 12.29% | 686 | 7.43% |
| Diabetes | 196 | 7.14% | 472 | 7.20% | 687 | 8.88% |
| CVD risk<sup>a</sup> | 196 | 0.30<br>(0.16) | 442 | 0.29<br>(0.18) | 684 | 0.28<br>(0.15) |
| Physically<br>active<sup>b</sup> | 191 | 60.21% | 418 | 30.38% | 627 | 52.95% |

*MMSE = Mini-Mental State Examination, ICV = intracranial volume, BMI = body-mass index, BP = blood pressure, CVD = cardiovascular disease risk, F = female. <sup>a</sup>Calculated based on the office-based version of the Framingham cardiovascular disease risk score<sup>38</sup>. <sup>b</sup>Regular engagement in moderate to vigorous physical activity (yes/no), defined as self-reported engagement > 3 hours (Betula) or > once per week (H70 and SNAC-K).*

### MRI acquisition and processing

MRI scanning was performed with a 3T General Electric scanner for Betula, a 3T Philips Achieva scanner for H70, and a 1.5T Philips Intera scanner for SNAC-K. For Betula, T1-weighted anatomical images were acquired with a 3D fast spoiled gradient echo sequence (TR: 8.2 ms, TE: 3.2 ms, field of view: 250 x 250 mm, flip angle: 12°, slice thickness: 1mm). For SNAC-K, T1-weighted anatomical images were acquired with a 3D MPRAGE sequence (TR: 15ms, TE: 7ms, field of view: 240 x 240 mm, flip angle: 15°, slice thickness: 1.5mm), and for H70, T1-weighed anatomical images were acquired with a 3D TFE SENSE sequence (TR: 7.2 ms, TE: 3.2 ms, field of view: 256 x 256 mm, flip angle: 9°, slice thickness: 1 mm).

In each cohort, volume of the lateral ventricles and the total intracranial volume (ICV) were estimated with FreeSurfer (Betula: version 7.1, SNAC-K: version 5.1; H70: version 7.2)^36^. For Betula and H70, MRI data for longitudinal analyses was processed with FreeSurfer’s longitudinal pipeline^37^, whereas for SNAC-K all data was processed with a cross-sectional pipeline. Ventricular volume was adjusted for ICV via the method of covariance (adjusted volume = raw volume – *b*(ICV – mean ICV), where *b* is the slope of the regression of the regional volume on ICV. In each cohort, slope and mean ICV were calculated from the baseline sample. For longitudinal analyses of Betula and H70, an overall estimate of ICV from FreeSurfer’s longitudinal pipeline for each individual was used to adjust raw ventricular volumes at all timepoints. For longitudinal analyses of SNAC-K, mean ICV across the available cross-sectional FreeSurfer ICV estimates was calculated for each individual and used for adjustment at all timepoints.

### Cognitive assessments

A score of general cognitive function was computed based on a principal component analysis (PCA) of four age-sensitive domains (see Supplementary material for details of cognitive assessments). For Betula, the cognitive domains included episodic memory, verbal fluency, visuospatial reasoning, and perceptual speed. For SNAC-K, the cognitive domains included episodic memory, verbal fluency, executive function, and perceptual speed. For H70, episodic memory, working memory, visuospatial reasoning, and perceptual speed were included. Outliers > 3.5 *SD*s from the mean on any cognitive measure were excluded prior to the PCA. The first principal component was extracted, explaining approximately half of the variance in each cohort (Betula: 48.83%, SNAC-K: 53.54%, H70: 47.70%, see Supplementary Table S2 for task loadings). PCA was computed on the baseline data and factor scores then estimated for all timepoints.

### Health and lifestyle factors

To examine predictors of brain maintenance, data on educational level (number of years), body-mass index (BMI), systolic blood pressure (BP, mmHg), current smoking status (yes/no), diagnosis of diabetes (yes/no), regular engagement in moderate to vigorous physical activity (yes/no, defined as self-reported engagement > 3 hours per week for Betula, and > once per week for SNAC-K and H70) were drawn from each cohort. Moreover, for each participant, total cardiovascular disease risk was estimated based on the office-based version of the Framingham risk score that considers age, sex, medication for hypertension, systolic BP, BMI, and smoking status^38^. Detailed description of the assessment of health and lifestyle measures in each cohort is available in the Supplementary material.

### Statistical analyses

Statistical analyses were conducted with R (version 4.3.2) and JASP (version 0.18.1). Linear mixed effects models were used to assess longitudinal changes in brain integrity and cognition, including a random effect of participant and fixed effects of time, baseline age, sex, and an interaction between time and baseline age. The interaction term was retained in the final model if it improved model fit, as indicated by a significant likelihood ratio test, and was not included for H70, which had an age-homogenous sample.

To investigate whether maintenance of global brain integrity was associated with better preserved cognitive function, we conducted two sets of subgroup analyses. We used a ventricular volume threshold of mean + 3 *SD*s estimated based on the younger sample in Betula to classify each older individual to high (ventricular volume ≤ mean + 3 *SD*s of young) or low (ventricular volume > mean + 3 *SD*s of young) brain integrity subgroup. Two different subgroup classifications were conducted: one considering only baseline MRI data and one considering MRI data at baseline and follow-up. For the longitudinal data, individuals were classified into the high brain integrity subgroup if they maintained ventricular volume within the younger reference sample (≤ mean + 3 *SD*s) at baseline and each follow-up timepoint where data was available (see Supplementary Figure S1 for visualization). In Betula, 121 individuals (61.73%) were classified into the high brain integrity group at baseline and 61 (50.83%) over the follow-up. In SNAC-K, 297 (62.92%) individuals were classified into the high-brain integrity group at baseline and 161 (54.76%) over the follow-up. In H70, 463 (67.30%) individuals were classified into the high brain integrity subgroup at baseline and 187 (45.95%) over the follow-up.

Baseline subgroup differences in cognition were assessed with ANCOVAs, controlling for age, sex, and study (pooled analyses). Longitudinal subgroup differences in cognition were assessed with linear mixed effects models, including a random effect of participant and fixed effects of baseline age, sex, study (pooled analyses), subgroup, and an interaction between subgroup and time. Linear mixed effects models were implemented with the R package lme4 (version 1.1-35), and *p*-values estimated via the Satterthwaite’s degrees of freedom method with the R package lmerTest (version 3.1-3). Lifestyle factors predicting brain maintenance (i.e., subgroup membership) were assessed with logistic regressions. We examined both regression models adjusted for study cohort only (referred to as unadjusted analyses), and models additionally adjusted for baseline age and sex. Birth cohort effects on baseline brain integrity were examined with ANOVAs, including individuals aged 70-72 years, who were born maximum 15 years apart across the cohorts (SNAC-K, *N* = 69, born 1929-1931, Betula, *N* = 46, born 1937-1940, H70, *N* = 688, born 1944). Alpha-level was set to *p* < 0.05 (two-tailed, uncorrected) for all analyses.

## Results

### Longitudinal changes in global brain integrity and general cognitive function

In each cohort, over follow-up time, ventricular volume increased (Betula, *β* = .20, 95% CI [.19, .22], *t* = 21.55, *p* < .001; SNAC-K, *β* = .20, 95% CI [.19, .22], *t* = 24.13, *p* < .001; H70, *β* = .30, 95% CI [.28, .32], *t* = 30.12, *p* < .001) and cognitive performance decreased (Betula, *β* = -.25, 95% CI [-.31, -.19], *t* = 8.68, *p* < .001; SNACK, *β* = -.06, 95% CI [-.10, -.03], *t* = 3.28, *p* = .001; H70, *β* = -.26, 95% CI [-.30, -.23], *t* = 13.59, *p* < .001) (see Figure 1). Accelerated increases in ventricular volume with higher baseline age were evident in Betula, *β_age x_* _time_ = .03, 95% CI [.01, .05], *t* = 2.96, *p* = .003, and SNAC-K, *β_age x time_* = .03, 95% CI [.02, .05], *t* = 3.78, *p* < .001, whereas for cognition, no evidence for an interaction between baseline age and time was observed in either cohort (*p*s > .272). Age and time interactions were not tested for H70, which had an age-homogeneous sample.

**Figure 1.**
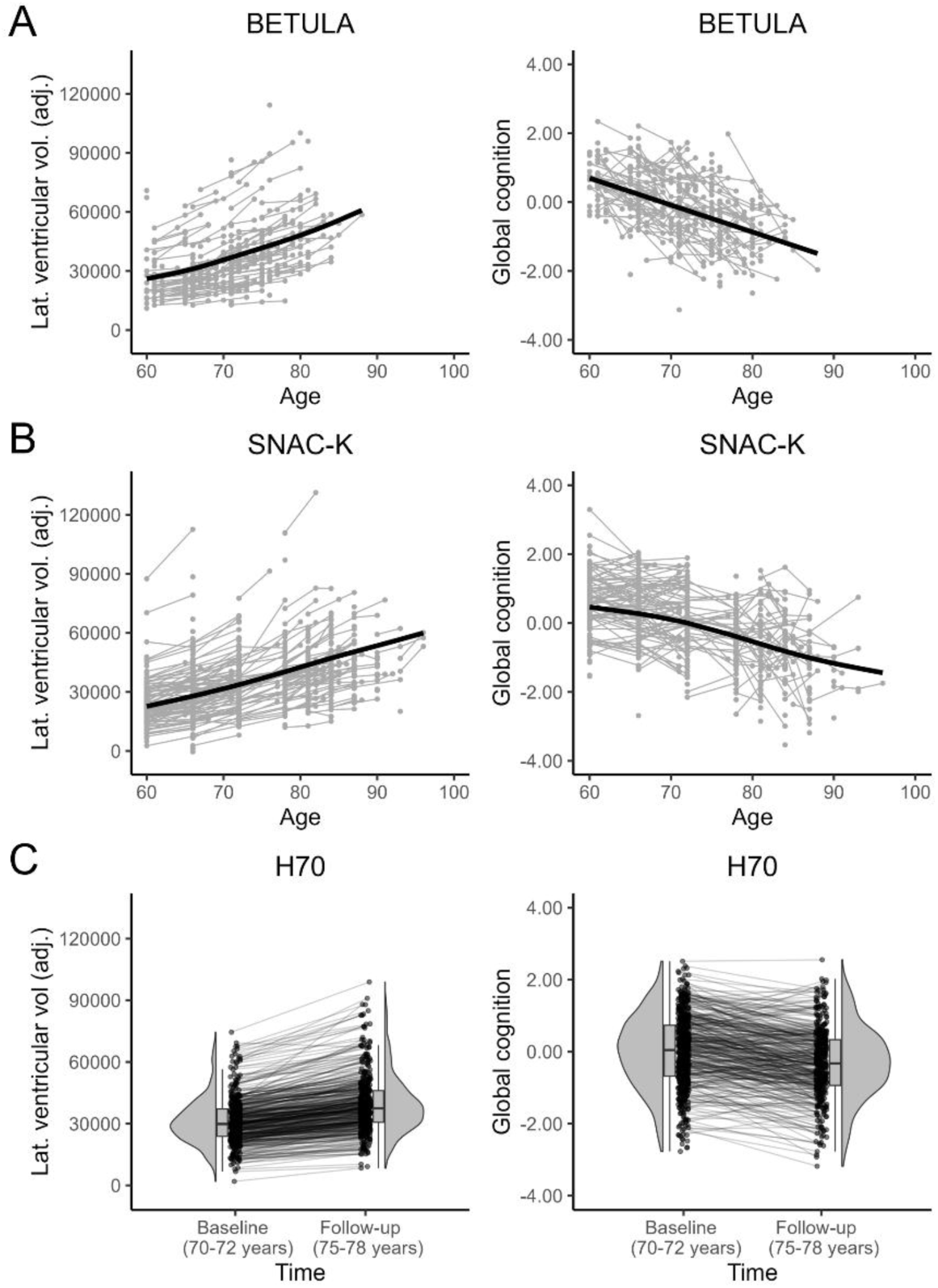
Trajectories of global brain integrity (indexed by volume of the lateral ventricles, adjusted for total intracranial volume) and global cognitive performance in each cohort (indexed by first principal component of four age-sensitive cognitive domains). Generalized additive mixed models were fitted to data from (A) Betula and (B) SNAC-K to visualize age-related trajectories (black line), using the R package gamm4.

### Brain maintenance is associated with better baseline cognition and less steep cognitive decline over time

To investigate whether maintenance of more youth-like brain integrity was associated with better preserved cognition, we performed two sets of subgroup analyses (see Methods for details). By definition, the subgroups formed considering baseline MRI data only differed in baseline ventricular volume (Betula, *t*(194) = 17.19, *p* < .001; SNAC-K, *t*(470) = 25.14, *p* < .001; H70, *t*(686) = 30.97, *p* < .001). Subgroups formed considering both baseline and follow-up MRI data differed in baseline ventricular volume (Betula, *β_subgroup_* = -1.21, 95% CI [-1.46, -0.96], *t* = 9.38, *p* < .001; SNAC-K, *β_subgroup_* = -1.19 95% CI [-1.36, -1.03]*, t* = 14.10, *p* < .001; H70, *β_subgroup_* = -1.16, 95% CI [-1.29, -1.02], *t* = 17.22, *p* < .001), and change in ventricular volume over time (Betula, *β_subgroup x time_* = -.15, 95% CI [-.18, -.12], *t* = 9.40, *p* < .001; SNAC-K, *β_subgroup x time_* = -.13, 95% CI [-.16, -.10], *t* = 8.48, *p* < .001; H70, *β_subgroup x time_* = -.18, 95% CI [-.22, -.15], *t* = 10.00, *p* < .001, see Supplementary material Figure S2).

Pooled analyses of baseline subgroup differences in cognition revealed that individuals with more youth-like brain integrity displayed better general cognitive function at baseline, *F*(1, 1196) = 20.84, *p* < .001, *η_p_^2^* = .02, controlling for age, sex, and study (see Figure 2A). This effect was also observed at the level of individual cohorts, with the high brain integrity subgroup demonstrating better cognitive performance in Betula, *F*(1, 192) = 4.94, *p* = .027, *η_p_^2^* = .03, SNAC-K, *F*(1, 409) = 5.54, *p* = .019, *η_p_^2^* = .01, and H70, *F*(1,589) = 9.76, *p* = .002, *η_p_^2^* = .02.

**Figure 2.**
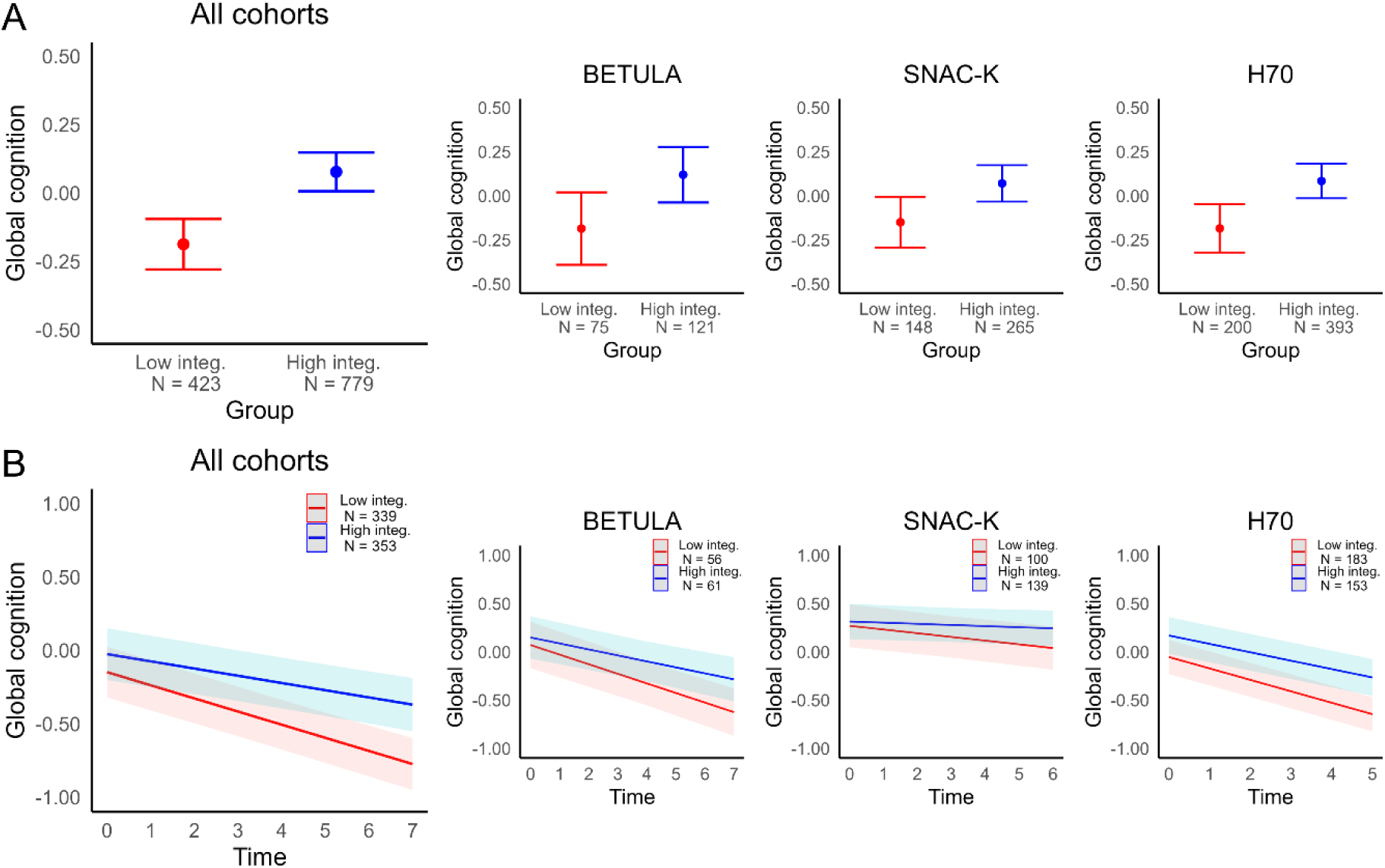
(A) Baseline differences in general cognitive function for high (i.e., ‘youth-like’) versus low (i.e., ‘aged’) brain integrity subgroups formed considering baseline MRI data. (B) Longitudinal changes in general cognitive function for high versus low brain integrity subgroups formed considering baseline and follow-up MRI data. Plots illustrate (A) estimated marginal means of cognitive performance from ANCOVAs, and (B) predicted cognitive performance from linear mixed effects models, controlling for baseline age, sex and study (pooled analyses).

Considering MRI data from both the baseline and follow-up, those who maintained more youth-like brain integrity also exhibited less steep cognitive decline over time (see Figure 2B). Pooled analyses of longitudinal subgroups indicated a significant interaction between subgroup and time for general cognitive function, *β* = .11, 95% CI [.06, .16], *t* = 4.32, *p* < .001, with a more pronounced effect of time on cognition observed in the low brain integrity subgroup, *β* = -.24, 95% CI [-.28, -.21], *t* = 13.29, *p* < .001, in comparison to the high brain integrity subgroup, *β* = -.14, 95% CI [-.17, -.10], *t* = 7.61, *p* < .001. A similar pattern was evident at the cohort level, with those who maintained global brain integrity over the follow-up displaying less steep cognitive decline in Betula, *β_subgroup x time_* = .12, 95% CI [.00, .23], *t* = 2.01, *p* = .046, and H70, *β_subgroup x time_* = .08, 95% CI [.00, .16], *t* = 2.06, *p* = .040, with a trend-level effect in the same direction observed for SNAC-K, *β_subgroup x time_* = .08, CI [.00, .15], *t* = 1.93, *p* = .055. For H70, the longitudinal subgroups also differed in cognitive performance at baseline, *β_subgroup_* = .23, CI [.02, .43], *t* = 2.16, *p* = .031.

Sensitivity analyses indicated consistent subgroup differences in cognition across varying ventricular volume thresholds used for the subgroup classification (see Supplementary Figure S3).

### Lifestyle factors predicting brain maintenance

We next sought to characterize potential health and lifestyle factors that promoted the maintenance of global brain integrity (Tables 2 and 3). Here, we focus on pooled analyses, with cohort-specific analyses reported in the Supplementary material (Tables S3-S8).

**Table 2.** Baseline subgroup differences in demographic, lifestyle, and health factors (low brain integrity, N = 475; high brain integrity, N = 881). Odds ratios (OR) and p-values from logistic regressions predicting subgroup membership, controlling for ^1^study or ^2^study, age and sex.

|  | Low integrity | High integrity | OR<sup>1</sup><br>[95% CI] | p-value<sup>1</sup> | OR<sup>2</sup><br>[95% CI] | p-value<sup>2</sup> |
| --- | --- | --- | --- | --- | --- | --- |
| Age | 72.42<br>(6.22) | 68.79<br>(5.22) | 0.51<br>[0.45, 0.58] | **< .001** |  |  |
| Sex | 56.42% F | 53.58% F | 0.89<br>[.71, 1.12] | .327 |  |  |
| Education | 12.44<br>(4.01) | 13.34<br>(4.18) | 1.24<br>[1.10, 1.39] | **< .001** | 1.08<br>[0.96, 1.22] | .209 |
| BMI | 26.21<br>(4.12) | 25.96<br>(3.95) | 0.94<br>[0.84, 1.06] | .314 | 0.96<br>[0.85, 1.08] | .475 |
| BP systolic | 142.63<br>(19.11) | 140.33<br>(18.52) | 0.89<br>[0.80, 1.00] | **.045** | 0.98<br>[0.87, 1.11] | .778 |
| Smoker | 8.23% | 9.77% | 1.23<br>[0.84, 1.86] | .299 | 0.90<br>[0.59, 1.38] | .615 |
| Diabetes | 12.84% | 5.45% | 0.39<br>[0.26, 0.57] | **<.001** | 0.39<br>[0.26, 0.59] | **<.001** |
| CVD risk<sup>a</sup> | 0.32<br>(0.17) | 0.27<br>(0.15) | 0.70<br>[0.62, 0.78] | **< .001** | 0.80<br>[0.68, 0.93] | **.004** |
| Physically active<sup>b</sup> | 42.79% | 48.39% | 1.26<br>[0.99, 1.61] | .062 | 1.20<br>[0.93, 1.55] | .159 |

BMI = body mass index, BP = blood pressure, CVD = cardiovascular disease, F = female.
<sup>a</sup>Calculated based on the office-based version of the Framingham cardiovascular disease risk score<sup>38</sup>. <sup>b</sup>Regular engagement in moderate to vigorous physical activity (yes/no), defined as self-reported engagement > 3 hours (Betula) or > once per week (H70 and SNAC-K).

**Table 3.** Longitudinal subgroup differences in demographic, lifestyle, and health factors (low brain integrity, N = 412; high brain integrity, N = 409). Odds ratios (OR) and p-values from logistic regressions predicting subgroup membership, controlling for ^1^study or ^2^study, age and sex.

|  | Low integrity | High integrity | OR<sup>1</sup><br>[95% CI] | p-value<sup>1</sup> | OR<sup>2</sup><br>[95% CI] | p-value<sup>2</sup> |
| --- | --- | --- | --- | --- | --- | --- |
| Age | 71.19<br>(5.43) | 68.12<br>(5.35) | 0.54<br>[0.46, 0.63] | **<.001** |  |  |
| Sex | 55.83% F | 56.23% F | 0.99<br>[0.75, 1.31] | .969 |  |  |
| Education | 13.07<br>(4.18) | 13.76<br>(4.13) | 1.20<br>[1.05, 1.39] | **.009** | 1.10<br>[0.95, 1.27] | .223 |
| BMI | 25.99<br>(4.01) | 25.76<br>(3.66) | 0.93<br>[0.81, 1.07] | .312 | 0.97<br>[0.84, 1.12] | .662 |
| BP systolic | 141.64<br>(18.64) | 138.71<br>(18.14) | 0.84<br>[0.73, 0.97] | **.016** | 0.93<br>[0.80, 1.08] | .341 |
| Smoker | 7.06% | 7.82% | 1.06<br>[0.63, 1.81] | .817 | 0.72<br>[0.41, 1.27] | .252 |
| Diabetes | 9.25% | 4.65% | 0.48<br>[0.27, 0.84] | **.012** | 0.53<br>[0.29, 0.94] | **.034** |
| CVD risk<sup>a</sup> | 0.29<br>(0.16) | 0.24<br>(0.14) | 0.68<br>[0.58, 0.78] | **<.001** | 0.79<br>[0.64, 0.96] | **.021** |
| Physically active<sup>b</sup> | 45.79% | 50.26% | 1.32<br>[0.98, 1.77] | .068 | 1.23<br>[0.91, 1.68] | .179 |

BMI = body mass index, BP = blood pressure, CVD = cardiovascular disease, F = female.
<sup>a</sup>Calculated based on the office-based version of the Framingham cardiovascular disease risk score<sup>38</sup>. <sup>b</sup>Regular engagement in moderate to vigorous physical activity (yes/no), defined as self-reported engagement > 3 hours (Betula) or > once per week (H70 and SNAC-K).

Membership of the high brain integrity subgroup at baseline was predicted by younger age, *β* = -.68, *SE* = .07, *p* < .001, *OR* = 0.51, 95% CI [0.45, 0.58], higher education, *β* = .21, *SE* = .06, *p* < .001, *OR* = 1.24, 95% CI [1.10, 1.39], lower systolic BP, *β* = -.12, *SE* = .06, *p* = .045, *OR* = 0.89, 95% CI [0.80, 1.00], absence of diabetes, *β* = -.95, *SE* = .20, *p* < .001, *OR* = 0.39, 95% CI [0.26, 0.57], and lower cardiovascular risk score, *β* = -.36, *SE* = .06, *p* < .001, *OR* = 0.70, 95% CI [0.62, 0.78], in unadjusted analyses. Controlling for age and sex, absence of diabetes, *β* = -.93, *SE* = .21, *p* < .001, *OR* = 0.39, 95% CI [0.26, 0.59], and lower cardiovascular risk score, *β* = -.23, *SE* = .08, *p* = .004, *OR* = 0.80, 95% CI [0.68, 0.93], remained as significant predictors of brain maintenance.

Similarly, membership of the high brain integrity subgroup over the follow-up was predicted by younger age, *β* = -.61, *SE* = .08, *p* < .001, *OR* = 0.54, 95% CI [0.46, 0.63], higher education level, *β* = .19, *SE* = .07, *p* = .009, *OR* = 1.20, 95% CI [1.05, 1.39], lower systolic blood pressure, *β* = -.17, *SE* = .07, *p* = .016, *OR* = 0.84, 95% CI [0.73, 0.97], absence of diabetes, *β* = -.73, *SE* = .29, *p* = .012, *OR* = 0.48, 95% CI [0.27, 0.84], and lower cardiovascular risk score, *β* = -.39, *SE* = .08, *p* < .001, *OR* = 0.68, 95% CI [0.58, 0.78], in unadjusted analyses. After controlling for baseline age and sex, absence of diabetes, *β* = - .64, *SE* = .30, *p* = .034, *OR* = 0.53, 95% CI [0.29, 0.94], and lower cardiovascular risk score, *β* = -.24, *SE* = .10, *p* = .021, *OR* = 0.79, 95% CI [0.64, 0.96], remained as significant predictors of longitudinal subgroup membership.

### Birth cohort effects on global brain integrity

Lastly, we assessed potential birth cohort effects on global brain integrity (Figure 3). For this purpose, baseline data were pooled from age-matched (70-72 years) individuals in each cohort, born maximum 15 years apart (born 1929-1944). No significant effect of birth cohort on adjusted ventricular volume was observed, *F*(2,800) = 2.44, *p* = .088. Given that birth cohort effects were reported for ICV in previous work^29^, we further examined potential cohort differences on ICV and raw ventricular volume, however, similarly found no significant effects (*p*s > .228).

**Figure 3.**
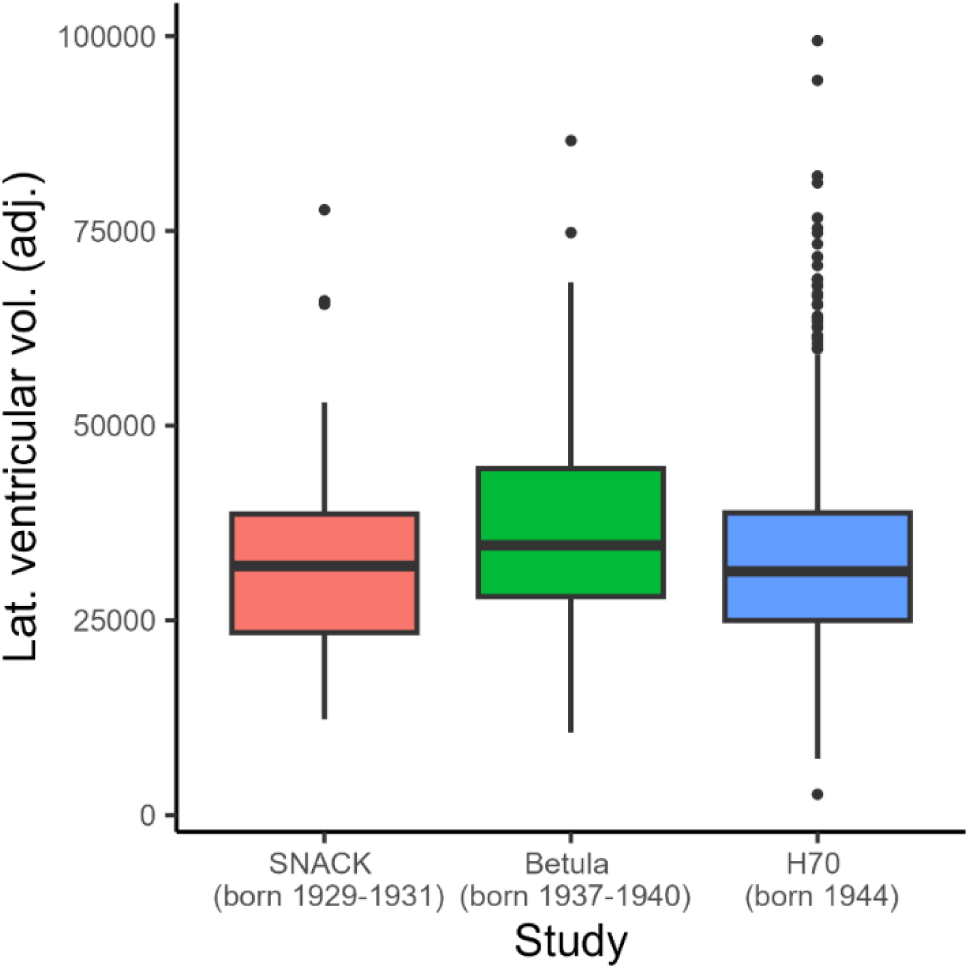
Baseline ventricular volume (adjusted for total intracranial volume) in 70-72-year-olds from each cohort (SNAC-K, *N* = 69, Betula, *N* = 46, H70, *N* = 688).

## Discussion

In this pooled analysis of three population-based Swedish cohorts, we provide evidence for a role of global brain maintenance in better preserved cognitive function in older age. Older individuals who maintained more youth-like global brain integrity displayed better general cognitive performance at baseline, and less longitudinal decline in cognition. Of lifestyle and demographic factors, the absence of diabetes and lower cardiovascular risk score were associated with higher likelihood of brain maintenance, highlighting the importance of managing cardiovascular and metabolic disease risk factors for promotion of brain and cognitive health in later life.

Previous accounts highlight brain maintenance (i.e., relative preservation of neural resources) as a critical determinant of individual differences in cognitive ageing^1^, however, large-scale longitudinal evidence for this proposal has been limited. Here, in pooled analyses of three population-based cohorts we show that those older individuals who maintained more youth-like global brain integrity also demonstrated better general cognitive performance at baseline, and less cognitive decline over a follow-up period of up to 7 years. This pattern was further largely replicated at the level of individual studies. Our findings align with previous studies demonstrating MRI markers of global brain integrity (e.g., total cortical atrophy, ventricular enlargement) to account for individual differences in cognitive ageing trajectories^4,39^. However, while we chose ventricular volume as our marker of global brain integrity in this multi-cohort analysis due to established links to cognitive decline^11,12^, good reliability and reproducibility^13,15,16^, we note that the reliance on a single metric means that we cannot evaluate the relative importance of *global* versus *local* brain maintenance in the current study.

Assessing lifestyle predictors of brain maintenance, we observed the absence of diabetes and lower cardiovascular disease risk to be associated with greater likelihood of brain maintenance at baseline and over the follow-up. Additionally, lower systolic blood pressure and higher education were associated with brain maintenance in unadjusted analyses, however, these associations did not survive controlling for age and sex. In addition to being associated with specific markers of vascular brain pathology (e.g., white matter hyperintensities)^26^, associations between cardiovascular risk burden and indices of overall brain structural integrity have been previously reported^25–27^. Our findings of a link between diabetes and lower likelihood of brain maintenance are further consistent with a recent analysis of the cross-sectional UK Biobank data, which observed diabetes to be among top lifestyle predictors of structural integrity of brain regions vulnerable to ageing^40^, as well as with meta-analytic evidence for accelerated brain ageing in individuals with type 2 diabetes^41^. In contrast to later life lifestyle factors, we did not observe significant birth cohort effects on ventricular volume when comparing 70-72-year-olds born maximum 15 years apart across the three cohorts. It is possible that larger differences in birth year may be required for detection of such trends, which have recently been reported across five decades of birth from the Framingham Heart Study^29^.

In the absence of longitudinal MRI data dating back to young adulthood, we used cross-sectional estimates of ventricular volume from a younger reference sample to classify older individuals as ‘youth-like’ or ‘aged’, akin to approaches previously applied for cognitive performance data^42^. While sensitivity analyses indicated consistent subgroup differences in cognition for varying ventricular volume thresholds estimated from the younger sample, it is important to acknowledge that longitudinal data is required to truly distinguish between those individuals who exhibit neural maintenance versus decline over midlife and older age. Moreover, we highlight that lifestyle and health predictors of brain maintenance were evaluated at baseline (i.e., age 60-93 years) in the current investigation, and we were therefore not able to assess potential temporal trends in the importance of lifestyle predictors over adulthood^26^.

To summarize, we provide evidence across three population-based cohorts that global brain maintenance is associated with better preserved general cognitive function in older age. Brain maintenance was predicted by absence of diabetes and lower overall cardiovascular disease risk, highlighting the importance of modifiable lifestyle factors for promotion of later life cognitive and brain health.

## Data availability

Access to cohort data used in this study can be requested via the Swedish National E-Infrastructure for Aging Research (https://www.near-aging.se/).

