## Supplementary material for "Global brain maintenance predicts well-preserved cognitive function: A pooled analysis of three longitudinal population-based Swedish cohorts"

**Assessment of cognitive performance in each cohort**

***Betula.*** Cognitive assessments for Betula have been described in detail in Nilsson et al. (1997, 2004). Episodic memory was assessed with five tasks at baseline and at the first follow-up, and with two tasks at the second follow-up. At each timepoint, summary score of the available episodic memory tasks was calculated. The two-item episodic memory summary score from the second follow-up was transformed into an estimate of the five-item summary score using a participant-specific scaling factor, as described previously (Johansson et al., 2020). The five episodic memory tasks completed at baseline and at the first follow-up consisted of immediate free recall of verb-noun sentences that were either verbally and visually presented or enacted by the participant (16 sentences per condition), delayed cued recall of nouns from the same sentences, and immediate free recall of 12 verbally presented nouns. At the second follow-up, only immediate free recall of sentences was administered (both visual/verbal and enacted condition). Verbal fluency was assessed by a category fluency task, where participants were instructed to produce as many professions starting with the letter B as they could within one minute. Visuospatial reasoning was assessed with a block design test (Wechsler, 1981). Perceptual speed was assessed with a letter digit substitution task, in which participants substitute digits with letters according to a transformation key as quickly as possible for the duration of one minute.

***SNAC-K.*** Cognitive assessments for SNAC-K have been described in detail in Laukka et al. (2013). Episodic memory was assessed with an immediate free recall of 16 verbally and visually presented nouns. Verbal fluency was assessed with two category fluency tasks, in which participants were asked to produce as many animals or professions as they could within one minute. Executive function was assessed with the Trail Making B test. Perceptual speed was assessed with a digit cancellation task (Zazzo, 1974) and a pattern comparison task (Salthouse & Babcock, 1991). For verbal fluency and perceptual speed, a summary score of the two available tasks was used.

***H70.*** Cognitive assessments for H70 have been described in detail in Rydberg Sterner et al. (2019). Episodic memory was assessed with a free recall of 12 objects shown to the participants (five minute delay), free immediate recall of 10 words read aloud by the participant, and free immediate recall of 10 garments verbally presented to the participant (BUS II, Buschke & Fuld, 1974). Working memory was assessed with the digit span forward and backward tasks (Wechsler, 1991), visuospatial reasoning with the block design test (Wechsler, 1981), and perceptual speed with a figure identification test (Dureman et al.,

1971). For episodic memory and working memory, a summary score of the individual tasks was used.

### Assessment of lifestyle and health factors in each cohort

For all cohorts, education was reported as years of formal education and body mass index (BMI) calculated as weight (kg) / height $(m)^2$. We further calculated an overall cardiovascular disease risk score for all participants using the office-based Framingham risk score function that considers age, sex, systolic blood pressure, medication for blood pressure, BMI, smoking status, and diabetes diagnosis (D'Agostino et al., 2008). This score represents the probability of occurrence of any cardiovascular disease event within a 10-year period.

***Betula.*** Systolic blood pressure (mmHg) was measured lying down, with the exception of eight participants for whom the measurements were taken while sitting. Current smoking status (yes/no) and diagnosis of diabetes (yes/no) were assessed via self-report. Medication for hypertension (yes/no) was assessed from self-reported use of prescription medications, considering antihypertensives (ATC code: C02), diuretics (ATC code: C03), beta blockers (ATC code: C07), calcium channel blockers (ATC code: C08) and renin-angiotensin system agents (ATC code: C09). Regular engagement in moderate to vigorous physical activity was assessed via self-report with the question "*How much time do you spend a typical week doing moderately strenuous activity that causes you to be warm (e.g., brisk walking, gardening, heavier housework, cycling, swimming*)?". Participants were categorized into regularly active (> 3h per week) or not (≤ 3h per week).

***SNAC-K.*** Systolic blood pressure (mmHg) was measured twice in a sitting position and the average of the two measurements calculated. Current smoking status (yes/no) was assessed via self-report. Diagnosis of diabetes (yes/no) was identified on the basis of a medical examination, antidiabetic drug use, diagnoses in the Swedish National Patient Register (ICD-9: code 250; ICD-10: code E11), or HbA1c ≥6.5% (48 mmol/mol) according to the American Diabetes Association criteria. Medication for hypertension (yes/no) was defined based on current use of antihypertensive agents (ATC codes: C02, C03, C07, C08, C09). Regular engagement in moderate to vigorous physical activity was evaluated via self-report with the question "*Have you regularly engaged in moderate to intense exercise in the last 12 months* (*e.g., jogging, long power walks, heavy-duty gardening, long bicycle rides, high-intensity aerobics, long distance ice skating, swimming, ball sports*)?". Participants were classified as regularly active (≥ several times per week) or not (≤ 2-3 times per month).

***H70.*** Systolic blood pressure (mmHg) was measured in a sitting position. Current smoking status (yes/no), diagnosis of diabetes (yes/no), and medication for hypertension (yes/no) were assessed via self-report. Engagement in regular moderate to vigorous physical activity was evaluated with the questions "*During the last 7 days, on how many days did you do vigorous physical activities like heavy lifting, heavy construction or gardening work, aerobics, running, or fast bicycling?*" and "*During the last 7 days, on how many days did you do moderate physical activities like bicycling, swimming, moderate construction or gardening work, or other at a moderate pace?".* The average number of days per week across the two questions was calculated and participants were classified as regularly active (> once per week) or not (≤ once per week).

Table S1. *Baseline characteristics of young to middle-aged participants (20-55 years old) in the Betula cohort (N = 60).*

| | Mean (SD) / Percentage |
|---|---|
| Age | 39.37 (9.41) |
| Sex | 50% F |
| Education | 14.85 (2.81) |
| MMSE | 28.55 (1.35) |
| Ventricular volume | 16902.86 (7567.65) |
| ICV | 1503106.04 (174058.58) |

*MMSE = Mini-Mental State Examination, ICV = intracranial volume, F = female.*

Table S2. *Loadings of each cognitive domain on the first principal component used to index general cognitive function in each cohort (Betula, N = 196; SNAC-K, N = 413; H70, N = 593).*

| Betula | | SNAC-K | | H70 | |
|---|---|---|---|---|---|
| Measure | Loading | Measure | Loading | Measure | Loading |
| EM | .75 | EM | .56 | EM | .59 |
| PS | .82 | PS | .82 | PS | .79 |
| FLU | .51 | FLU | .75 | VIS | .76 |
| VIS | .68 | EXC | .78 | WM | .60 |

EM = episodic memory, PS = perceptual speed, FLU = verbal fluency, VIS = visuospatial reasoning, EXC = executive function, WM = working memory.

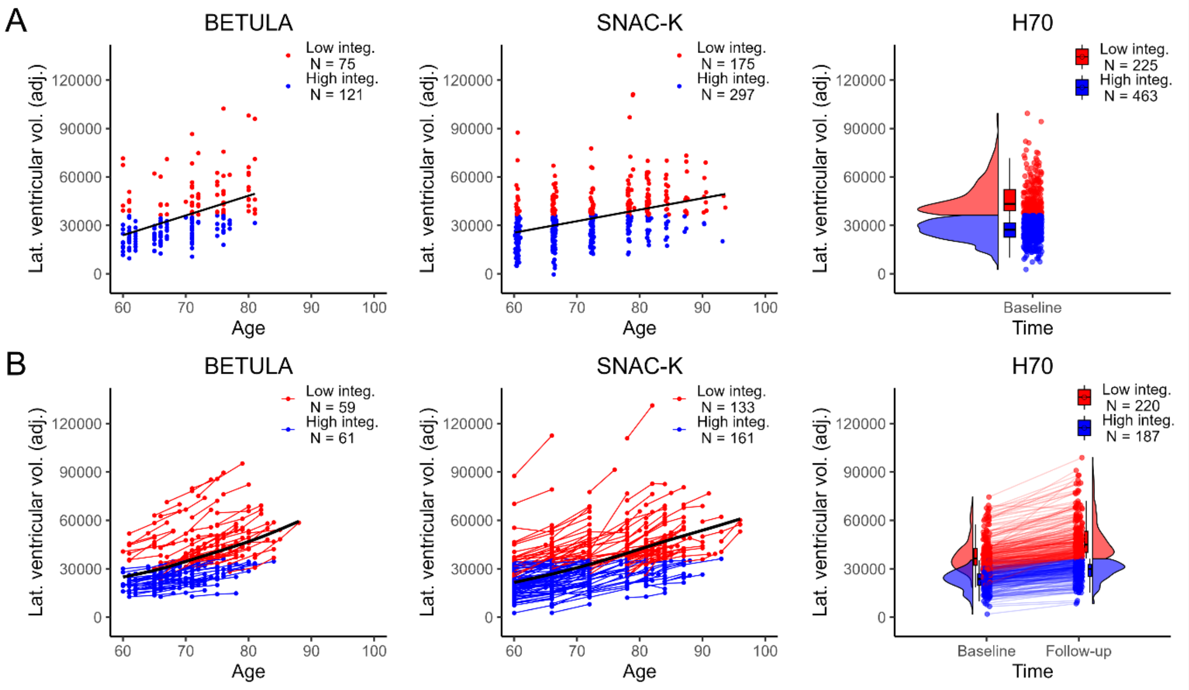


*Figure S1.* Classification of older individuals into subgroups of high (ventricular volume ≤ mean + 3SDs of the younger reference sample) or low (ventricular volume > mean + 3SDs of the younger reference sample) at A) baseline and B) over the entire follow-up period in each cohort.

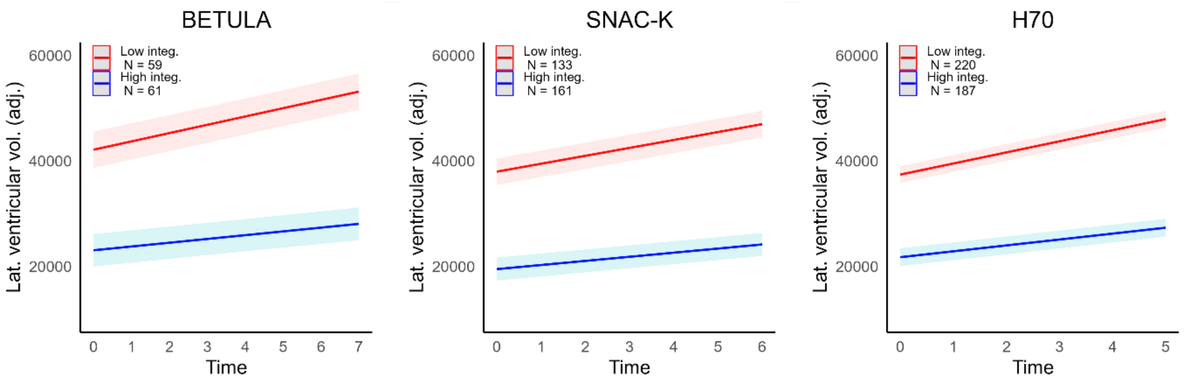


*Figure S2.* Longitudinal changes in ventricular volume in the subgroups defined based on baseline and follow-up MRI data. Plots illustrate predicted volume of the lateral ventricles (adjusted for total intracranial volume) from linear mixed effects models including a random effect of participant, and fixed effects of time, sex, baseline age, and subgroup, as well as an interaction between time and subgroup.

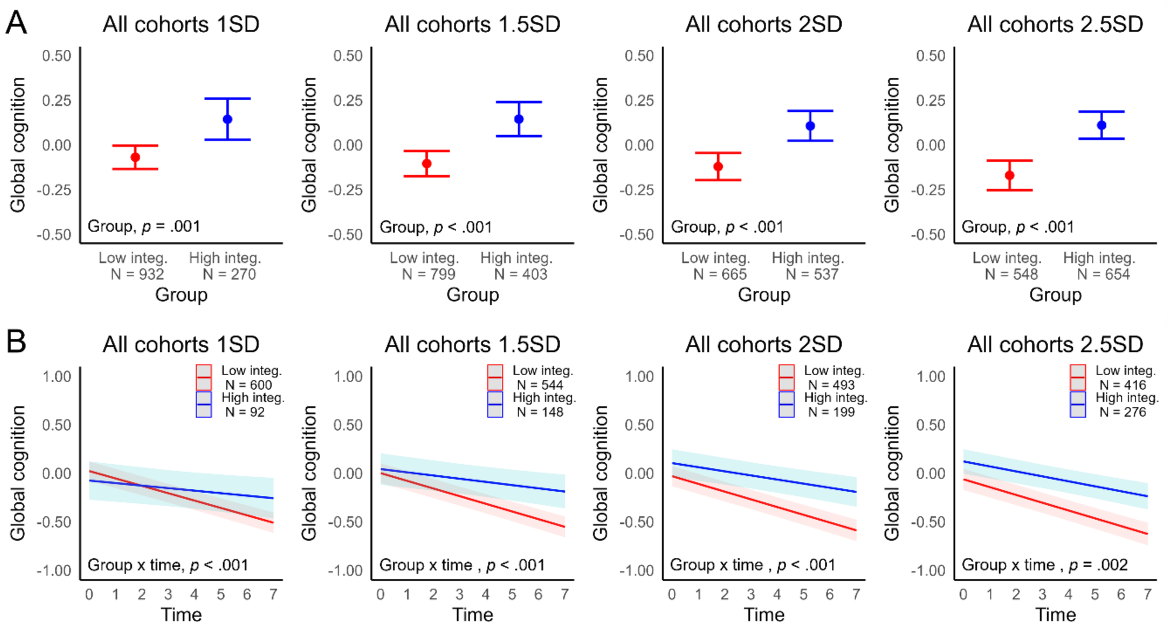


*Figure S3.* Sensitivity analyses of subgroup differences in A) baseline cognition and B) longitudinal changes in cognition for varying ventricular volume thresholds based on the Betula younger adult sample (mean + 1*SD* – mean + 2.5 *SD*s). Plots illustrate (A) estimated marginal means of cognitive performance from ANCOVAs, and (B) predicted cognitive performance from linear mixed effects models, controlling for baseline age, sex and study.

Table S3. *Baseline subgroup differences in demographic, health, and lifestyle factors in the Betula cohort (low integrity, N = 75; high integrity, N = 121). Odds ratios (OR) and p-values from logistic regression predicting subgroup membership, [1]unadjusted for covariates, or [2]adjusted for age and sex.*

| | Low integrity | High integrity | OR[1] [95% CI] | p-value[1] | OR[2] [95% CI] | p-value[2] |
|---|---|---|---|---|---|---|
| Age | 72.45 (6.35) | 66.92 (5.41) | 0.37 [0.26, 0.52] | **< .001** | | |
| Sex | 45.33% F | 51.24% F | 1.27 [0.71, 2.27] | .422 | | |
| Education | 12.04 (3.99) | 12.82 (4.39) | 1.20 [0.90, 1.63] | .217 | 0.71 [0.49, 1.01] | .059 |
| BMI | 26.42 (3.44) | 26.48 (3.66) | 1.02 [0.76, 1.36] | .918 | 1.06 [0.77, 1.46] | .723 |
| BP systolic | 147.03 (17.59) | 139.36 (17.05) | 0.64 [0.47, 0.86] | **.004** | 0.81 [0.56, 1.14] | .239 |
| Smoker | 8.00% | 8.26% | 1.04 [0.37, 3.16] | .948 | 0.38 [0.12, 1.30] | .111 |
| Diabetes | 14.67% | 2.48% | 0.15 [0.03, 0.49] | **.004** | 0.16 [0.03, 0.62] | **.013** |
| CVD risk | 0.37 (0.17) | 0.25 (0.14) | 0.45 [0.32, 0.62] | **< .001** | 0.60 [0.38, 0.93] | **.026** |
| Physically active | 61.64% | 59.32% | 0.91 [0.50, 1.65] | .750 | 1.00 [0.51, 1.94] | .991 |

*BMI = body mass index, BP = blood pressure, CVD = cardiovascular disease, F = female.*

Table S4. *Longitudinal subgroup differences (low integrity, N = 59; high integrity, N = 61) in demographic, health, and lifestyle factors in the Betula cohort. Odds ratios (OR) and p-values from logistic regression predicting subgroup membership, [1]unadjusted for covariates, [2]adjusted for baseline age and sex.*

| | Low integrity | High integrity | OR[1] [95% CI] | p-value[1] | OR[2] [95% CI] | p-value[2] |
|---|---|---|---|---|---|---|
| Age | 70.22 (5.52) | 65.56 (4.91) | 0.39 [0.25, 0.59] | **< .001** | | |

| | | | | | | |
|---|---|---|---|---|---|---|
| Sex | 49.15% F | 45.90% F | 0.88 [0.43, 1.80] | .722 | | |
| Education | 12.59 (4.34) | 13.07 (4.27) | 1.12 [0.78, 1.62] | .541 | 0.81 [0.53, 1.23] | .327 |
| BMI | 26.74 (3.78) | 26.45 (2.99) | 0.92 [0.64, 1.31] | .637 | 0.94 [0.63, 1.38] | .738 |
| BP systolic | 142.85 (16.64) | 135.66 (17.16) | 0.64 [0.43, 0.93] | **.025** | 0.81 [0.52, 1.22] | .308 |
| Smoker | 6.78% | 8.20% | 1.23 [0.31, 5.19] | .769 | 0.58 [0.13, 2.60] | .455 |
| Diabetes | 10.17% | 3.28% | 0.30 [0.04, 1.36] | .150 | 0.41 [0.05, 2.16] | .322 |
| CVD risk | 0.32 (0.16) | 0.24 (0.14) | 0.55 [0.36, 0.81] | **.004** | 0.73 [0.41, 1.26] | .260 |
| Physically active | 46.43% | 65.00% | 2.14 [1.02, 4.57] | **.045** | 2.17 [0.96, 5.00] | .065 |

*BMI = body mass index, BP = blood pressure, CVD = cardiovascular disease, F = female.*

Table S5. *Baseline subgroup differences in demographic, health, and lifestyle factors in the SNAC-K cohort (low integrity, N = 175; high integrity, N = 297). Odds ratios (OR) and p-values from logistic regression predicting subgroup membership, [1]unadjusted for covariates, or [2]adjusted for age and sex.*

| | Low integrity | High integrity | OR[1] [95% CI] | p-value[1] | OR[2] [95% CI] | p-value[2] |
|---|---|---|---|---|---|---|
| Age | 75.19 (8.60) | 67.35 (7.89) | 0.38 [0.31, 0.48] | **<.001** | | |
| Sex | 66.29% F | 54.21% F | 0.60 [0.41, 0.88] | **.010** | | |
| Education | 11.77 (3.74) | 13.23 (4.15) | 1.45 [1.20, 1.77] | **< .001** | 1.11 [0.89, 1.38] | .342 |
| BMI | 26.22 (4.26) | 26.04 (3.43) | 0.95 [0.79, 1.15] | .615 | 0.99 [0.80, 1.21] | .888 |
| BP systolic | 145.00 (19.68) | 141.17 (18.81) | 0.82 [0.67, 0.99] | **.041** | 0.96 [0.77, 1.19] | .716 |

| | | | | | | |
|---|---|---|---|---|---|---|
| Smoker | 7.43% | 15.15% | 2.23 [1.20, 4.41] | **.016** | 1.52 [0.76, 3.23] | .256 |
| Diabetes | 9.71% | 5.72% | 0.56 [0.28, 1.14] | .109 | 0.56 [0.26, 1.23] | .150 |
| CVD risk | 0.34 (0.18) | 0.26 (0.17) | 0.63 [0.51, 0.77] | **< .001** | 0.85 [0.61, 1.17] | .313 |
| Physically active | 26.67% | 32.46% | 1.32 [0.85, 2.07] | .217 | 1.03 [0.63, 1.69] | .896 |

*BMI = body mass index, BP = blood pressure, CVD = cardiovascular disease, F = female.*

Table S6. *Longitudinal subgroup differences in demographic, health, and lifestyle factors in the SNAC-K cohort (low integrity, N = 133; high integrity, N = 161). Odds ratios (OR) and p-values from logistic regression predicting subgroup membership, [1]unadjusted for covariates, or [2]adjusted for baseline age and sex.*

| | Low integrity | High integrity | OR[1] [95% CI] | p-value[1] | OR[2] [95% CI] | p-value[2] |
|---|---|---|---|---|---|---|
| Age | 73.20 (8.48) | 66.63 (7.34) | 0.43 [0.32, 0.55] | **< .001** | | |
| Sex | 63.16% F | 61.49% F | 0.93 [0.58, 1.50] | .769 | | |
| Education | 12.47 (3.96) | 13.59 (4.07) | 1.33 [1.05, 1.69] | **.019** | 1.13 [0.86, 1.47] | .382 |
| BMI | 26.33 (4.12) | 25.84 (3.46) | 0.88 [0.69, 1.11] | .269 | 0.96 [0.74, 1.23] | .739 |
| BP systolic | 145.69 (20.54) | 138.92 (18.19) | 0.70 [0.54, 0.89] | **.004** | 0.83 [0.64, 1.09] | .183 |
| Smoker | 8.27% | 11.80% | 1.48 [0.69, 3.34] | .322 | 0.82 [0.36, 1.98] | .657 |
| Diabetes | 9.02% | 4.35% | 0.46 [0.17, 1.17] | .112 | 0.53 [0.18, 1.46] | .227 |
| CVD risk | 0.32 (0.17) | 0.23 (0.15) | 0.53 [0.40, 0.68] | **< .001** | 0.73 [0.49, 1.08] | .122 |
| Physically active | 28.81% | 34.46% | 1.30 [0.77, 2.20] | .327 | 1.07 [0.60, 1.90] | .825 |

*BMI = body mass index, BP = blood pressure, CVD = cardiovascular disease, F = female.*

Table S7. *Baseline subgroup differences in demographic, health, and lifestyle factors in the H70 cohort (low integrity, N = 225; high integrity, N = 463). Odds ratios (OR) and p-values from logistic regression predicting subgroup membership, [1]unadjusted for covariates, [2]adjusted for age and sex.*

| | Low integrity | High integrity | OR[1] [95% CI] | p-value[1] | OR[2] [95% CI] | p-value[2] |
|---|---|---|---|---|---|---|
| Age | 70.26 (0.46) | 70.21 (0.43) | 0.90 [0.77, 1.05] | .179 | | |
| Sex | 52.44% F | 53.78% F | 1.06 [0.77, 1.45] | .742 | | |
| Education | 13.10 (4.13) | 13.54 (4.14) | 1.12 [0.95, 1.31] | .186 | 1.11 [0.95, 1.31] | .204 |
| BMI | 26.14 (4.23) | 25.79 (4.31) | 0.92 [0.79, 1.08] | .312 | .93 [0.79, 1.09] | .356 |
| BP systolic | 139.36 (18.68) | 140.07 (18.73) | 1.04 [0.89, 1.22] | .641 | 1.03 [0.88, 1.21] | .704 |
| Smoker | 8.93% | 6.71% | 0.73 [0.41, 1.34] | .300 | 0.74 [0.42, 1.36] | .321 |
| Diabetes | 14.67% | 6.06% | 0.38 [0.22, 0.64] | **< .001** | 0.37 [0.22, 0.64] | **< .001** |
| CVD risk | 0.29 (0.15) | 0.27 (0.14) | 0.85 [0.73, 1.00] | **.048** | 0.82 [0.68, 0.99] | **.043** |
| Physically active | 47.83% | 55.48% | 1.36 [0.97, 1.90] | .072 | 1.36 [0.98, 1.91] | .069 |

*BMI = body mass index, BP = blood pressure, CVD = cardiovascular disease, F = female.*

Table S8. *Longitudinal subgroup differences in demographic, health, and lifestyle factors in the H70 cohort (low integrity, N = 220; high Integrity = 187). Odds ratios (OR) and p-values from logistic regression predicting subgroup membership, [1]unadjusted for covariates, or [2]adjusted for baseline age and sex.*

| | Low integrity | High integrity | OR[1] [95% CI] | p-value[1] | OR[2] [95% CI] | p-value[2] |
|---|---|---|---|---|---|---|
| Age | 70.23 (0.43) | 70.24 (0.45) | 1.02 [0.84, 1.24] | .841 | | |

| | | | | | | |
|---|---|---|---|---|---|---|
| Sex | 53.18% F | 55.08% F | 1.08 [0.73, 1.60] | .702 | | |
| Education | 13.55 (4.22) | 14.14 (4.12) | 1.15 [0.95, 1.40] | .161 | 1.15 [0.95, 1.41] | .153 |
| BMI | 25.58 (3.97) | 25.48 (3.99) | 0.97 [0.80, 1.18] | .784 | 0.97 [0.80, 1.19] | .796 |
| BP systolic | 138.92 (17.56) | 139.53 (18.39) | 1.03 [0.85, 1.26] | .734 | 1.03 [0.85, 1.26] | .750 |
| Smoker | 6.39% | 4.28% | 0.65 [0.26, 1.56] | .351 | 0.65 [0.26, 1.56] | .349 |
| Diabetes | 9.13% | 5.35% | 0.56 [0.25, 1.21] | .151 | 0.57 [0.25, 1.22] | .156 |
| CVD risk | 0.27 (0.14) | 0.25 (0.12) | 0.87 [0.71, 1.06] | .178 | 0.84 [0.65, 1.08] | .177 |
| Physically active | 55.34% | 58.62% | 1.14 [0.76, 1.72] | .520 | 1.15 [0.77, 1.73] | .496 |

*BMI = body mass index, BP = blood pressure, CVD = cardiovascular disease, F = female.*
